# Circumventing drug treatment? Polyethylenimine functionalized nanoparticles inherently and selectively kill patient-derived glioblastoma stem cells

**DOI:** 10.1101/2020.12.14.422717

**Authors:** Neeraj Prabhakar, Joni Merisaari, Vadim Le Joncour, Markus Peurla, Didem Şen Karaman, Eudald Casals, Pirjo Laakkonen, Jukka Westermarck, Jessica M. Rosenholm

**Affiliations:** Pharmaceutical Sciences Laboratory, Faculty of Science and Engineering, Åbo Akademi University, Turku, 20520, Finland; Turku Bioscience Centre, University of Turku and Åbo Akademi University, Turku, 20520 Finland; Institute of Biomedicine, Faculty of Medicine, University of Turku, Turku, 20520, Finland; Translational Cancer Medicine Research Program, Faculty of Medicine, University of Helsinki, Helsinki, 00014, Finland; Department of Biomedical Engineering, Faculty of Engineering and Architecture, İzmir Kâtip Çelebi University. İzmir, 35620, Turkey; School of Biotechnology and Health Sciences, Wuyi University, Jiangmen 529020, China; Laboratory Animal Center, HiLIFE – Helsinki Institute of Life Science, University of Helsinki, Helsinki, 00014, Finland

## Abstract

Glioblastoma (GB) is the most frequent malignant tumor originating from the central nervous system. Despite breakthroughs in treatment modalities for other cancer types, GB remains largely irremediable due to its high degree of intratumoral heterogeneity, infiltrative growth, and intrinsic resistance towards multiple treatments. These resistant and aggressive sub-populations of GBs including the glioblastoma stem cells (GSCs) can circumvent treatment. GSCs act as a reservoir of cancer-initiating cells; they are a major challenge for successful therapy. We have discovered, as opposed to well-reported anti-cancer drug based therapeutical approach for GB therapy, the role of polyethylenimine (PEI) in inducing selective death of patient-derived GSCs via lysosomal membrane rupturing. Even at very low doses (1 μg/ml), PEI surface-functionalized mesoporous silica nanoparticles (PEI-MSNs), without any additional anti-cancer drug, very potently and selectively killed multiple GSC lines. Very importantly, PEI-MSNs did not affect the survival of well-established GB cells, or other type of cancer cells even at 25x higher doses. Remarkably, any sign of predominant cell death pathways such as apoptosis and autophagy was absent. Instead, as a potential explanation for their GSC selective killing function, we demonstrate that the internalized PEI-MSNs accumulated inside the lysosomes, subsequently causing a rupture of the vulnerable lysosomal membranes, exclusively in GSCs. As a further evaluation, we observed blood-brain-barrier (BBB) permeability of these PEI-MSNs *in vitro* and *in vivo*. Taking together the recent indications for the vulnerability of GSCs for lysosomal targeting, and GSC selectivity of the PEI-MSNs described here, the results suggest that PEI-functionalized nanoparticles could have a potential role in the eradication of GSCs.

## Introduction

Glioblastoma (GB) is the most common, aggressive, and lethal form of primary brain tumors in adults[1,2]. The prognosis of patients affected by GB remains limited with a median survival of approximately 12-18 months[3]. The current clinical practices for patient treatments include surgery, chemo- and radiotherapy. The treatments are challenged by major complications because of the highly invasive nature of GB cells, intratumoral heterogeneity, and the intrinsic resistance of GB cells towards therapies[4]. It has been reported that even after surgery and chemo- and radiotherapy, glioblastoma cells invade neighboring normal brain leading to currently uncurable recurrence in patients[5]. Current therapeutic approaches leave the resistant and aggressive sub-populations including GSCs untreated[6]. In the case of GB, GSCs belong to a small subgroup of cells with a phenotypically distinctive character: the ability and unlimited potential to differentiate, self-renew, and form new tumors. GSCs are one of the main causes of resistance, recurrence, and mortality in GB, thus, novel therapeutic approaches are needed to target the GSC population[7].

Conventional targeting of the stem cell populations within brain tumors has shown some success at the preclinical level[8]. The strategies used to target stem cells have mainly focused on signaling pathways[9] and dendritic cell-based immunotherapy[10]. Another potential strategy for treatment involves the application of PARP (poly (ADP-ribose) polymerase) inhibitors. The PARP inhibitor (ABT-888) enhances apoptosis when used in combination with TMZ (temozolomide) and radiation[11]. GSCs are known to survive harsh conditions, such as lack of oxygen (hypoxia) and nutrients[12]. GSCs can modify their metabolic machinery to enhance the glucose uptake via the high-affinity glucose transporter GLUT3. Therefore, GLUT3 receptors have shown to be a potential therapeutic target for glioblastoma GSCs[13]. Recently, Lucki *et al.* reported a drug-like small molecule cRIP GBM that selectively induces apoptosis in GSCs[14]. Another recent report by Le Joncour *et al.* showed that invasive GB cells display vulnerable lysosomes, and that lysosomal membrane permeabilization could be achieved by application of a cationic amphiphilic antihistamine class drug, clemastine (Tavegil™), a first-generation histamine H1 antagonist [15]. To develop more selective therapies, a deeper understanding of the intracellular biology and microenvironment of GSCs would be vital.

During recent years, silica-based nanoparticles have gained vast attention in therapy, diagnosis, and theranostics[16]; albeit limited examples exist specifically for brain cancer treatment. A few promising studies have been published to date [17–22], suggesting that their advantageous characteristics should be exploitable also in this therapeutic area. The most prominent advantage of mesoporous silica nanoparticles (MSNs) is perhaps their drug delivery capability, being able to efficiently carry high payloads of poorly soluble drugs; equipped with controlled release functions, ability to cross biological barriers e.g. the cell membrane, and in the best case scenario, delivering the drug in a targeted fashion[23–29]. The silica surface is inherently negatively charged, which in general does not maximize attraction to the negatively charged cellular membranes. Thus, hyperbranched polyethylenimine (PEI), a polycationic polymer with a high amount of amino groups[30] is widely applied for enhancing the cellular uptake of nanoparticles to achieve efficient delivery of therapeutic payloads to cells [31,32]. In addition to enhancing cellular uptake, PEI is widely believed to promote endosomal escape via the proton-sponge effect[33,34]. The proton-sponge hypothesis suggests that cationically surface-functionalized nanoparticles allow endo/lysosomal swelling by intake of water molecules, eventually leading to the disintegration of endo/lysosomal membranes[35,36]. Given that nanoparticles naturally accumulate in lysosomes upon cellular internalization as a result of endocytic mechanisms, lysosomal membrane disruption could be a potential route to exploit the vulnerability of GSCs.

In the present study, we set out to evaluate the *in vitro* and *in vivo* potential of PEI surface-functionalized MSNs (PEI-MSNs) as drug carriers to treat GB using patient-derived GSCs (BT-3-CD133^+^, BT-12, BT-13 cells)[15]. Nevertheless - and most remarkably - without carrying any drugs, PEI-MSNs were able to induce selective cell death of GSCs but not established GB cells without stem cell characteristics. MSNs without PEI coating did also not induce this effect, which led us to postulate that the selective cell death may have occurred via rupturing of the vulnerable lysosomal membrane. Subsequently, we performed in-depth intracellular microscopic analysis on BT-12 cells to understand the role of PEI functionalization in cell death. The results obtained by confocal and TEM imaging predominantly suggest the involvement of the “proton-sponge mechanism” induced by PEI-MSNs, leading to rupture of the lysosomal membrane. Additionally, to deduce the potential of this mechanism to be exploited in a therapeutic setting, we showed successful penetration of PEI-MSNs through BBB models both *in vitro* and *in vivo*. Furthermore, the *in vitro* blood-brain tumor-barrier (BBTB) model[37] confirmed accumulation of PEI-MSNs in the lysosomes of BT-12 GSC. Potentially, this discovery of the inherent role of PEI-MSNs in selectively eradicating otherwise highly resistant GSCs presents a novel vulnerability to exploit for brain cancer (GB) treatment.

## Materials and Methods

Unless otherwise noted, all reagent-grade chemicals were used as received, and Millipore water was used in the preparation of all aqueous solutions. Cetylmethylammonium bromide (CTAB, AR) was purchased from Fluka. 1,3,5-Trimethyl-benzene (TMB,99%) was purchased from ACROS. Decane (99%) was purchased from Alfa Aesar. Anhydrous toluene (AR), ethylene glycol (AR), tetraethyl orthosilicate (TEOS, AR), 3-aminopropyltriethoxysilane (APTES, AR), NH4OH (30 wt%, AR), were purchased from Sigma Aldrich. Aziridine was used in the preparation for hyperbranched surface modification of MSNs and purchased from Menadiona S.L.Pol. Industrial company.

### Preparation and characterization of hyperbranched PEI functionalized mesoporous silica nanoparticles (PEI-MSNs)

Mesoporous silica nanoparticles were prepared according to a protocol from our previously published work[38]. The MSNs were prepared by co-condensation of TEOS and APTES as silica sources. Briefly, a mixed solution was prepared by dissolving and heating CTAB (0.45 g) in a mixture of DI water (150 mL) and ethylene glycol (30 mL) at 70 °C in a reflux-coupled round flask reactor. Ammonium hydroxide (30 wt%, 2.5 mL) was introduced to the reaction solution as the base catalyst before TEOS (1.5 mL) and APTES (0.3 mL) was added to initiate the reaction. Decane (2,1 mL) and TMB (0.51 mL) were used as swelling agents before the addition of the silica sources, decane was added 30 min before TMB and after the addition of TBM, the synthesis solution was mixed for 1.5 h. The molar ratio of used reagents in the synthesis of MSN was 1TEOS : 0.19APTES : 0.18CTAB : 0.55TMB : 1.6 decane : 5.9NH_3_ : 88.5 ethylene glycol : 1249H_2_O. For inherent fluorophore labeling of the MSNs, TRITC was pre-reacted with APTES in a molar ratio of (APTES:TRITC) 3:1 in ethanol (0.5 mL) under vacuum for 2 h. Subsequently, the pre-reaction solution was added to the synthesis solution before the addition of TEOS. The reaction was allowed to proceed for 3 h at 70 °C. Then, the heating was stopped where after the as-synthesized colloidal suspension was aged at 70 °C without stirring for 24 h. After the suspension was cooled to room temperature, the suspension was separated by centrifugation. After collecting the particle precipitate, the template removal was carried out by the ion-exchange method. Briefly, the collected particles were extracted three times in ethanolic NH_4_NO_3_ solution, washed with ethanol[18], and resuspended in DMF for long-term storage. The surface modification of MSNs with hyperbranched PEI by surface-initiated polymerization was carried out according to an in-house-established protocol[30]. To initiate PEI polymerization from the MSNs surfaces, aziridine was used as a monomer with toluene as solvent, in which the MSN substrate was suspended in the presence of catalytic amounts of acetic acid. The suspension was refluxed under atmospheric pressure overnight at RT, filtered, washed with toluene, and dried under vacuum at 313 K. Henceforth, the obtained nanoparticles are abbreviated as PEI-MSNs. Full redispersibility of dried, extracted, and surface-functionalized MSN was confirmed by redispersion of dry particles in HEPES buffer at pH 7.2 and subsequent dynamic light scattering (DLS) measurements (Malvern ZetaSizer NanoZS). The fine architecture of the nanoparticles was further confirmed by transmission electron microscopy (Jeol JEM-1200EX electron microscope) operated at 80 kV. The success of surface polymerization was confirmed by zeta potential measurements (Malvern ZetaSizer NanoZS).

### Cell culture

Established human glioblastoma cell line T98G (VTT Technical Research Centre, Turku, Finland in 2010) was cultured in Eagle MEM (Sigma-Aldrich), supplemented with 10% heat-inactivated FBS (Biowest), 2 mM L-glutamine and penicillin (50 U/mL)/streptomycin (50 μg/mL). The patient-derived GSCs BT-3-CD133^+^, BT-12 and BT-13[7] were cultured in Dulbecco’s modified Eagle’s medium with Nutrient Mixture F-12 (DMEM/F12, Gibco) supplemented with 2 mM L-glutamine, 2% B27-supplement (Gibco), 50 U/ml penicillin and 50 μg/mL streptomycin, 0.01 μg/mL recombinant human fibroblast growth factor-basic (FGF-b, Peprotech), 0.02 μg/mL recombinant human epidermal growth factor (EGF, Peprotech) and 15 mM HEPES-buffer. The blood-brain-tumor barriers were established as previously described[26]. Mouse endothelial cells from brain microvessels (bEND3) were maintained in DMEM (Lonza) supplemented with 10% decomplemented FBS (Lonza), 2 mM L-glutamine (Sigma) and penicillin:streptomycin (50 U/mL and 50 μg/mL respectively). Mouse immortalized astrocytes (HIFko) were maintained in Basal Eagle Medium 1 (BME-1, Sigma) supplemented with 5% decomplemented FBS (Lonza), 1 M HEPES (Sigma), 2 mM L-glutamine (Sigma) 100 mM sodium pyruvate (Sigma), 3 g D-glucose and penicillin:streptomycin (50 U/mL and 50 μg/mL respectively). All cell lines were kept in a humidified atmosphere of 5% CO_2_ at 37 °C. For colony growth and microscopy GSC populations were cultured as monolayers on Matrigel (Becton Dickinson) coated dishes.

### Western blotting and antibodies

BT-12 cells were treated with 10 μg/mL PEI-MSNs for 24h and 48h. They were lysed in 2x Laemmly buffer (4% SDS, 20% glycerol, 120mM Tris) and resolved by SDS-PAGE gel (BioRad, Country). Proteins were transferred to nitrocellulose membranes (Bio-Rad). Membranes were blocked with 5% milk-TBS and incubated with a required dilution of primary and 1:5000 dilution of secondary antibody in 5% Milk-TBS-Tween 20 for a required duration of time and visualized with Odyssey (LI-COR Biosciences, Nebraska, USA). The membrane was blocked using 5% milk in Tris-buffered saline (TBS) and incubated with a primary antibody PARP-1 (sc-7150, 1:1000) and P62 (sc-28359, 1:500 dilution) from Santa Cruz Biotechnology. Antibodies for cPARP (ab32064) (1:1000 dilution) was acquired from Abcam, LC3-β (2775s) (1:1000) from Cell Signaling. Loading control antibodies for β-actin (sc-47778) (1:10,000 dilution) was from Santa-Cruz Biotechnology. Secondary antibodies were purchased from LI-COR, mouse (926-32212), and rabbit (926-68021).

### Colony formation assay

An optimized number of cells (3 × 10^3^ to 10 × 10^3^) were seeded in 24-well plates (Sigma-Aldrich) and allowed to attach. After 24 hours cells were treated with 1-50 μg/mL of PEI-MSNs. After 72h medium was replaced with fresh medium and the cells were incubated for another 72h or until the control well was confluent. Cell colonies were fixed with methanol dilutions and stained with 0.2% crystal violet (CV) solution in 10% ethanol for 15 min at room temperature. Plates were dried and scanned with Epson Perfection V700 Photo scanner. Quantifications were performed with ImageJ by using the Colony area plugin[39]. Data were normalized and presented as a percent of the control.

### Light microscopy

#### Immunofluorescence (Early endosomes and Lysosomes)

BT-12 GSCs were grown on as monolayers on Matrigel (Becton Dickinson) coated glass coverslips. BT-12 GSCs were treated with 10 μg/mL of PEI-MSNs conjugated with TRITC (Tetramethylrhodamine-isothiocyanate) for 48h. BT-12 GSCs were fixed with 4% PFA (Paraformaldehyde) for 10 min. The cells were permeabilized using 0.1% Triton X-100 for 10 min and blocked with horse serum. The 1° anti-EEA1 (goat) antibody for recognition of early endosomes (Santa Cruz Biotechnology, USA) was prepared (1:100) in PBS (10% horse serum). The 1° anti-LAMP-1 (mouse) antibody for recognition of lysosomes (Abcam, UK) was prepared (1:100) in PBS (10% horse serum). Antibody incubation was performed overnight at +4 °C. The cells were washed three times with PBS; Alexa 488 secondary (Anti-goat and anti-mouse) antibodies (Sigma-Aldrich, US) in PBS were added to the cells at RT for 1h. The cells were mounted on coverslips using VECTASHIELD (4′,6-diamidino-2-phenylindole). The microscopy setup consisted of Zeiss 780 (Zeiss, Germany) confocal microscope, PMT, and 100X oil objective. DAPI was excited by 405 lasers and emission was collected in the blue channel. Alexa 488 (early endosomes and lysosomes) was excited with 488nm argon laser and emission was collected by green channel (510-550 nm). The TRITC labeled PEI-MSNs were excited by 561 nm laser and emission were collected (575-610 nm).

#### Mitochondrial staining

BT-12 GSCs were grown on as monolayers on Matrigel (Becton Dickinson) coated glass coverslips and further, treated with 10 μg/mL of PEI-MSNs conjugated to FITC (Fluorescein isothiocyanate) for 48h. Cell medium (0.5 mL) was collected from the plate and mixed with 0.2 μL of Mitotracker Orange^®^ (Thermo Fisher Scientific Inc, USA) returned to the cells drop-by-drop. The cells were finally incubated for 20 min at 37 °C. The cells were washed 3x with PBS, fixed for 10 min with 4% PFA, and mounted using VECTASHIELD (4′,6-diamidino-2-phenylindole) on glass slides for microscopy. The microscopy setup consisted of Zeiss 780 (Zeiss, Germany) confocal microscope, PMT, and 100X oil objective. DAPI was excited by 405 lasers and emission was collected in the blue channel. FITC-conjugated PEI-MSNs were excited with 488 nm argon laser and emission was collected by green channel (510-550 nm). The Mitotracker Orange^®^ was excited by 561 nm laser and emission were collected at 575-610 nm.

### Transmission electron microscopy (TEM)

BT-12 GSCs were grown on as monolayers on Matrigel (Becton Dickinson) coated glass coverslips glass coverslips and further, treated with 10 μg/mL of PEI-MSNs 24 and 72h. The BT-12 GSCs were fixed with 5% glutaraldehyde s-collidine buffer, post-fixed with 2% OsO_4_ containing 3% potassium ferrocyanide, dehydrated with ethanol, and flat embedded in a 45359 Fluka Epoxy Embedding Medium kit. Thin sections were cut using an ultramicrotome to a thickness of 100 nm. The sections were stained using uranyl acetate and lead citrate to enable detection with TEM. The sections were examined using a JEOL JEM-1400 Plus transmission electron microscope operated at 80 kV acceleration voltage[40].

### *In vitro* Blood-Brain Tumor Barrier

Murine blood-brain-barriers in a dish (BBB) were established according to a previously published protocol[26]. Briefly, mouse brain microvessel endothelial cells (bEND3) were co-cultured in Transwell inserts with immortalized mouse astrocytes (HIFko). After 6 days, BBB dishes were placed on BT-12 organoids on glass coverslips to complete the blood-brain tumor-barriers (BBTB) and 100 ng of PEI-MSI were added on the endothelial side. After 24h, BBTB dishes were stained with LysoTracker Red DND-99 according to the manufacturer’s recommendations (Invitrogen) before fixation with ice-cold 4% PFA (10 min) and nuclear counterstaining with DAPI (1 μg/mL, Sigma). BT-12 coverslips and Transwell membranes containing both bEND3 and HIFko cells were cut and mounted on Mowiol 4-88 (Sigma) and imaged on a Zeiss LMS880 confocal microscope.

To quantify the cell viability of the bEND3, astrocytes, and BT-12 cells from the BBTB, cells were gently detached with accutase (Sigma) collected, counted and 5×10^5^ cells/mL were transferred in a 96-well plate. 10 μL of 3-(4,5-Dimethylthiazol-2-yl)-2,5-Diphenyltetrazolium Bromide (MTT;5 mg/ml in PBS) was added on the cells before incubating for 2h at 37°C. Eventually, cells were lysed 10% SDS, 10 mM HCl) o/n and the absorbance was measured at 540 nm using Multiskan Ascent software version 2.6 (Thermo Labsystems). Results were expressed as the % of absorbance relative to the control, untreated BBTB cells.

### *In vivo* procedures

Intracranial implantation of U87MG-GFP or BT-12 cells was performed as previously described[15]. Briefly, 8-week old female NMRI:Rj nude mice were implanted with 10^5^ cells in 10 μL in the right striatum. After 20 days of tumor growth, 100 μg of PEI-MSN in PBS were injected in the caudal vein (100 μL) or intranasally (3 dosages of 5 μL given every two hours). After 8h, animals were euthanized and brains were snap-frozen in −50°C isopentane (Honeywell). Brain cryosections (9 μm) were cut using a cryotome (ThermoFisher), collected on Superfrost Ultra slides (ThermoFisher), and fixed in a ice-cold 4% PFA bath. Brain microvessels were stained overnight using a rat anti-mouse PECAM-1/CD31 (1:400, 553370, BD Pharmingen). Cell nuclei were counterstained with DAPI (1 μg/mL, Sigma), samples were mounted with Mowiol 4-88, and imaged on a Zeiss LMS880 confocal microscope.

## Results

Examination of hydrodynamic size and ζ-potential values of PEI-functionalized MSNs in HEPES buffer solution (25 mM, pH 7.2) at the concertation of 0.1 mg/mL yielded a hydrodynamic mean size of 124 ± 12 nm with a low polydispersity index (PDI) value of 0.09, indicating a monodispersed colloidal suspension of PEI-MSNs. In addition, the ζ-potential value of PEI-MSNs in HEPES buffer (+39±4 mV) ascertained the high net positive charge on MSN surfaces owing to successful surface modification with PEI. The mesoscopic ordering of the MSN structure before the surface modification was examined by transmission electron microscopy (TEM) imaging of the samples **(Fig. S1)**. As presented in the TEM micrographs, spherical particles with an approximate size of 50 nm with a porous structure were obtained for dried powder.

### PEI-MSNs exhibit specific toxicity towards GSCs

The PEI-MSNs were applied (1-50 μg/mL) to T98G (established GB cell line), BT-3-CD133^+^, BT-12, and BT-13 (patient-derived GSCs) cells (**Fig. S2**) and the colony formation was followed by crystal violet staining **(Fig. 1A)**. The efficiency of colony formation was quantified by using the “ColonyArea” ImageJ plugin[39] **(Fig. 1B)**. The exposure of PEI-MSNs to GSC cells resulted in pronounced inhibition of colony growth even at particle concentration as low as 1 μg/mL. However, no significant effect on the growth of T98G, A172, and U87MG (GB) cells or MDA-MB-231 breast carcinoma and HeLa cervical carcinoma cells were observed even at 50 μg/mL PEI-MSN concentration **(Fig. 1, S3 and S5**).

**Figure 1.**
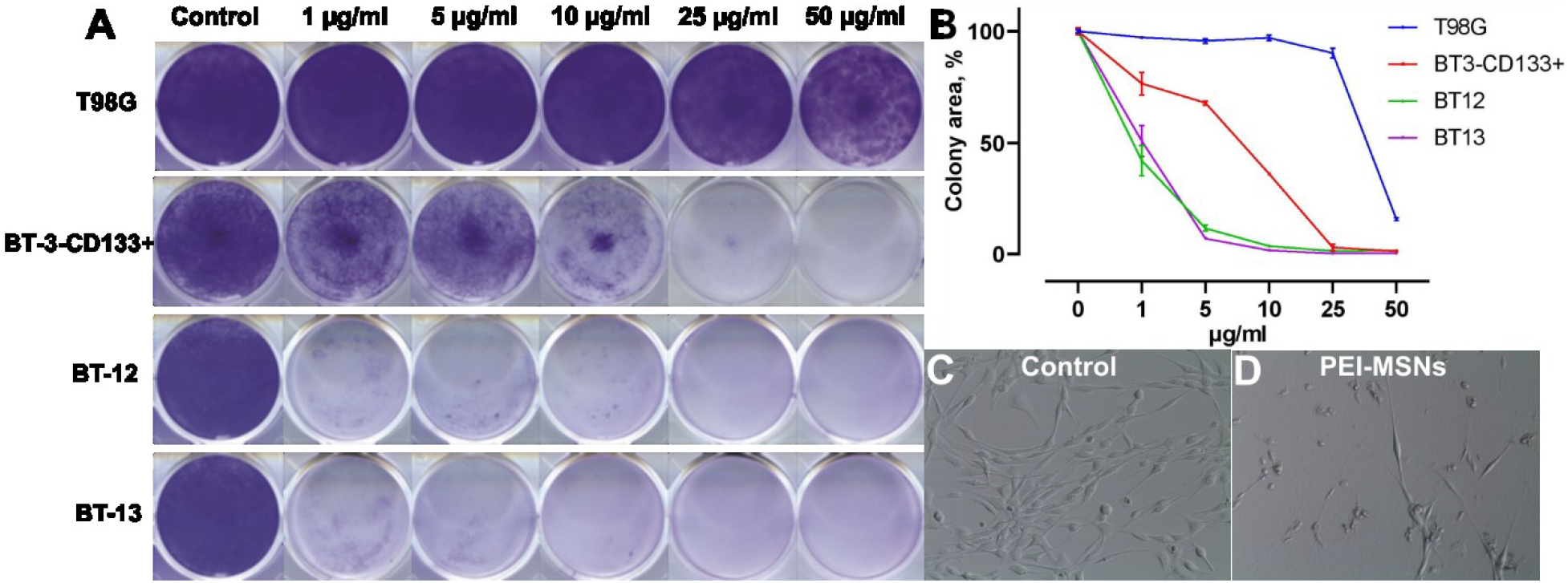
Selective death of patient-derived GSCs induced by PEI-MSNs. A) Colony growth assay of T98G (GB cell line), BT-3-CD133^+^, BT-12, and BT-13 (GSCs) cells treated with 1-50 μg/mL of PEI-MSNs. B) Quantification of colony growth by using the “ColonyArea”. C-D) Representative images of BT-3-CD133^+^ cells without (C) or with (D) PEI-MSN treatment.

At high PEI-MSN concentration (50μg/ml) also the T98G, U87MG, and A172 (GB) showed reduced colony growth in comparison to the control-treated cells (**Fig. 1B, S3 and S5**). These observations can be correlated to the well-reported fact that PEI can induce non-specific toxicity to cells if applied at higher concentrations[41–45]. Importantly, we further verified that MSNs without PEI did not cause cytotoxicity at the concentration range of 1-50 μg/mL **(Fig. S4)**. Thus, our results show that PEI functionalization played a very critical role in the induction of selective death of patient-derived GSCs, especially at low (1-5 μg/mL) concentrations.

### GSCs show no induction of apoptosis or autophagy after PEI-MSN treatment

We further investigated the role of PEI-MSNs in the induction of selective death of patient-derived GSCs. The BT-12 and BT-13 GSCs were the most sensitive to even low (1-5 μg/mL) PEI-MSN concentrations. Based on these results, we then selected the BT-12 GSCs for an in-depth analysis. Initially, the possible roles of autophagy or apoptosis in GSCs cell death were investigated by studying the cleavage of PARP-1 as an apoptosis marker in cells treated with PEI-MSNs [46–48]. PEI-MSN treatment of BT-12 GSCs for 24h and 48h did not affect the expression levels of this apoptotic biomarker (Fig. 2A).

**Figure 2.**
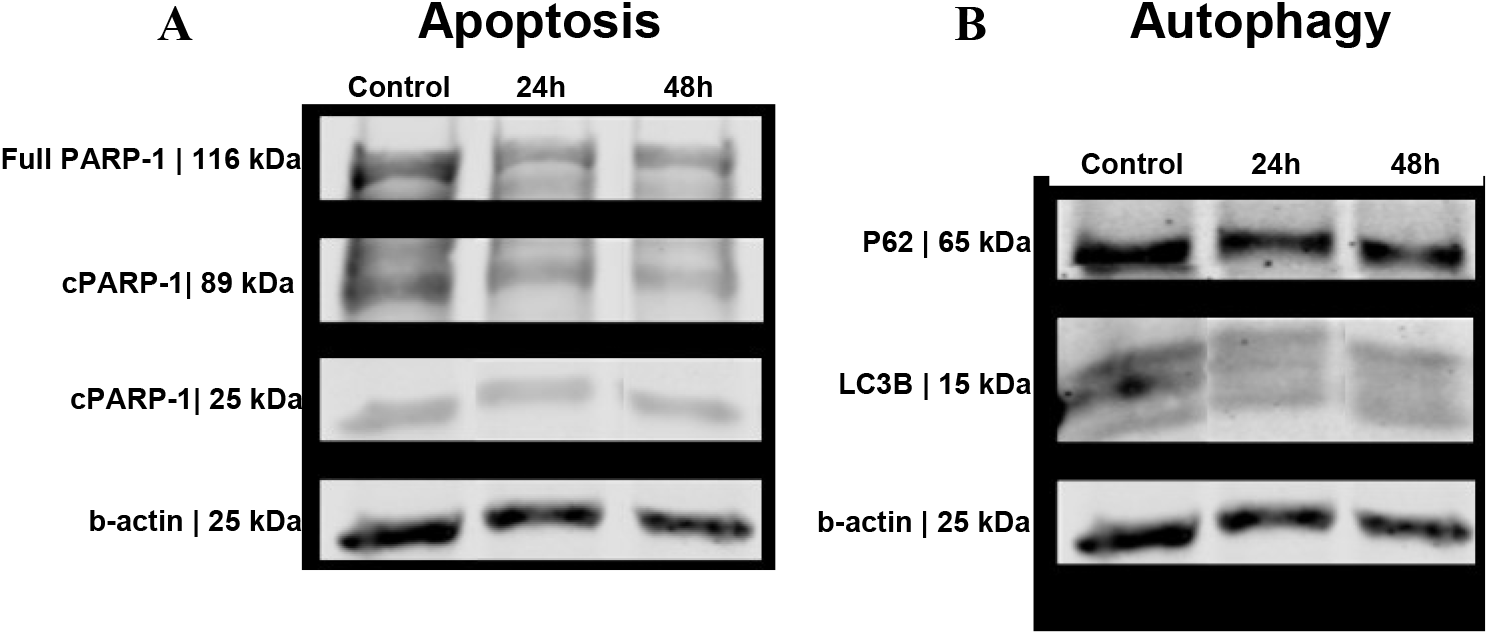
Western blot analysis of apoptosis and autophagy biomarkers in BT-12 GSCs treated with 10 μg/mL PEI-MSNs for 24h and 48h. A) Expression levels of apoptotic biomarkers of the full-length PARP-1, PARP-1C, and cPARP. B) Expression levels autophagy biomarkers P62 and LC3B.

To further determine the possible effect on autophagy, we studied the expression of specific autophagy biomarkers P62 and LC3B[49–52] in PEI-MSN treated cells. The biomarker expression of PEI-MSN treated cells was very similar to that of control cells, and no significant increase in autophagy-related biomarkers was observed **(Fig. 2B).** Thereby, these results suggest that cell death was not mediated by apoptosis or autophagy.

### PEI-MSNs localize within the cytoplasmic space and lysosomes

To understand the potential cell death mechanism of BT-12 GSCs, we studied the intracellular localization of PEI-MSNs by confocal microscopy. We selected early endosomes (EEA1), nucleus (DAPI), mitochondria (Mitotracker), and lysosomes (LAMP-1) to comprehend the PEI-MSNs interactions with the intracellular organelles in BT-12 GSCs **(Fig. 3)**. **Fig. 3** shows that upon 48h treatment, PEI-MSNs were mostly co-localized with the lysosomal marker (LAMP-1) in the BT-12 GSCs.

**Figure 3.**
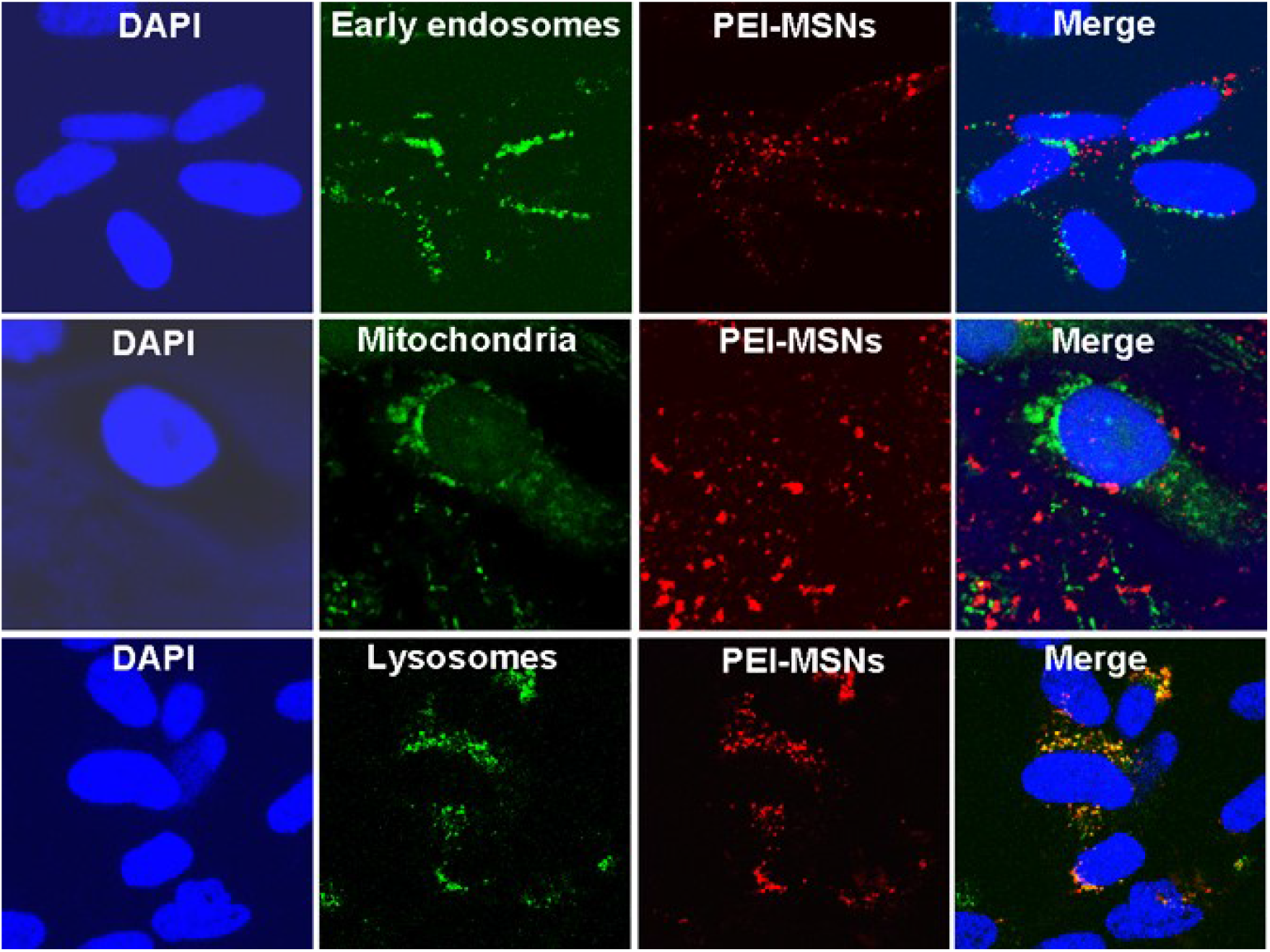
Localization of the PEI-MSNs in the treated BT-12 GSCs by confocal microscopy. The intracellular localization of PEI-MSNs (red) was studied using markers of early endosomes, mitochondria, and lysosomes (green color). The nuclei were visualized by using DAPI (blue). The co-localization of PEI-MSNs with lysosomes is seen in yellow.

PEI-MSNs did not localize within the nucleus. Besides, a non-significant amount of PEI-MSNs overlapped with either early endosomes (EEA1) or mitochondria. Individual PEI-MSNs (50 nm in diameter) were beyond the limit of resolution by confocal microscopy[53]. Despite that, the co-localization of the PEI-MSNs with the lysosomal marker (LAMP-1) was very frequently observed **(Fig. 3 and Fig. S6).** This was anticipated, given that nanoparticles typically enter cells by endocytosis after which they are transported to lysosomes[40,54,55]. Detection of the endosomal escape of PEI-MSNs is also beyond the detection limit of confocal microscopy and thus, this analysis cannot be used to detect possible “proton sponge effect” via membrane destabilization. Therefore, intracellular localization studies with confocal microscopy alone were not adequate to understand the full picture of intracellular interactions within the BT-12 GSCs. Therefore, subsequent TEM imaging was performed to investigate how PEI-MSNs interacts at the sub-cellular level.

### PEI-MSNs cause membrane rupture of lysosomes in GSCs leading to cell death

TEM imaging of the treated BT-12 GSCs revealed the widespread dissemination of PEI-MSNs **(Fig. 4A-D)** throughout the cytoplasmic space of cells. Moreover, we also observed that three distinct types of vesicular accumulation of PEI-MSN in the BT-12 GSCs: 1) vesicles filled with PEI-MSNs **(Fig. 4A)**, 2) empty vesicles with PEI-MSNs localized in the proximity to a vesicular membrane **(Fig. 4B)**, and 3) vesicles semi-filled with PEI-MSNs **(Fig. 4C-D)**. In summary, PEI-MSNs appeared localized in vesicles, mostly accumulated; as well as PEI-MSNs in non-vesicular spaces mostly as individual PEI-MSN particles **(Fig. 4)**.

**Figure 4.**
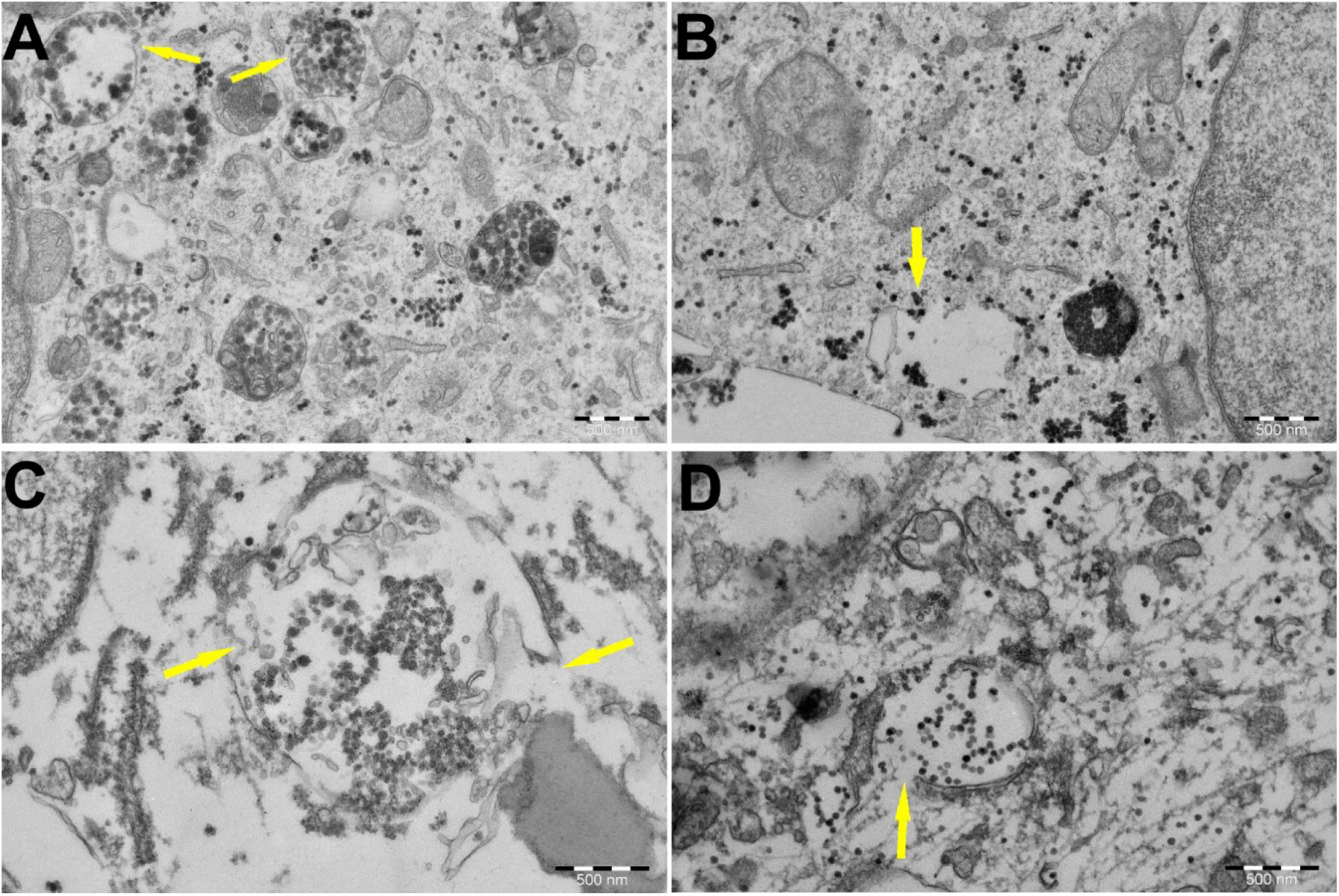
TEM imaging of BT-12 GSCs revealed the subcellular localization of the PEI-MSNs. PEI-MSNs were localized throughout the cells. A) Yellow arrows mark the potential endosomal escape of PEI-MSNs via vesicular membrane rupture. B) PEI-MSNs in close-proximity of a vesicle. C-D) A close-up view of affected vesicles suggesting a rupture of membranes and widespread localization of PEI-MSNs.

TEM imaging of the vesicle membranes (yellow arrows in Fig. **4A-D**) indicates the permeabilization and potential escape of PEI-MSNs from these damaged vesicles.

### PEI-MSNs cause morphological abnormalities in GSCs

Finally, we also observed morphological abnormalities **(Fig. 5A-B)** and structural damage of mitochondria in the PEI-MSN-treated BT-12 GSCs **(Fig. 5C-D)**. We did not observe PEI-MSNs permeating the nuclear space **(Fig. S7A-B)**. The easily recognizable abnormalities observed in the ultrastructure included: prevalent cytoplasmic localization of PEI-MSNs throughout the treated cells, loss of vesicle integrity, mitochondrial swelling, and rupture of cristae in comparison to the BT-12 control cells **(Fig. S8A-D)**. Theoretically, the process of endosomal trafficking begins with the early endosomes. The endosomal payload can be either recycled to the plasma membrane via recycling endosomes or it can advance to the late endosomes and lysosomes for degradation[56–58]. The proton sponge hypothesis indicates that PEI functionalization promotes escape from the endolysosomal pathway through rupture of the membrane. Numerous studies propose that membrane permeabilization occurs in the lysosomes[59,60]. Lysosomal membrane destabilization can lead to a triggered discharge of lysosomal enzymes to the cytoplasm, which eventually can cause cell death[61,62]. In the case of BT-12 GSCs, evidence from the confocal and TEM imaging suggests that the lysosomal membrane disruption by the PEI-MSNs could be a potential mechanism for BT-12 GSCs cell death.

**Figure 5.**
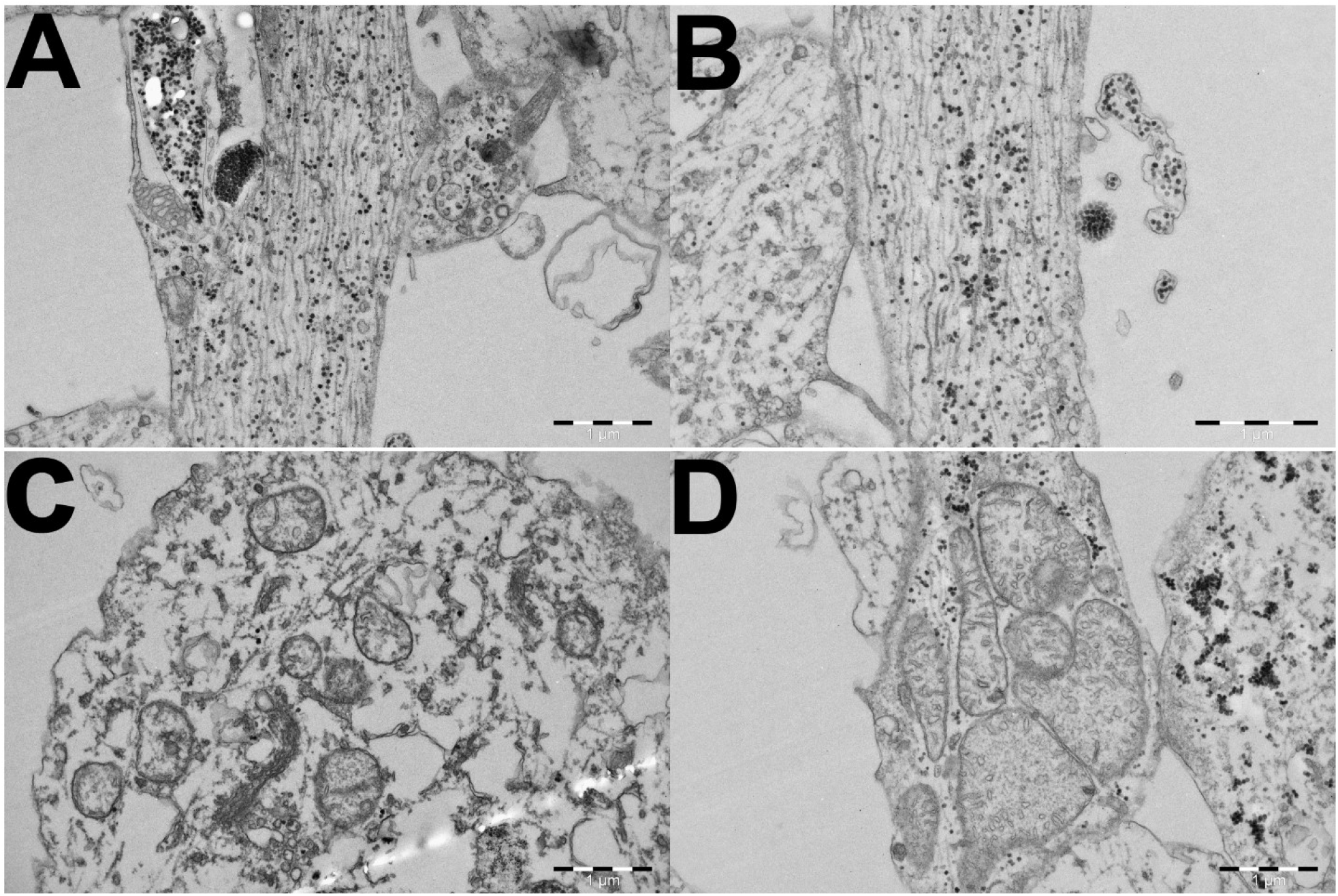
The cellular fate of PEI-MSN in the treated BT-12 GSCs. A-B) Endosomally escaped PEI-MSNs can be observed widespread within the cells, tandem with loss of structural integrity of the cells was detected. C-D) Abnormalities in the mitochondrial morphology, mitochondrial swelling, and rupture of the cristae.

### PEI-MSNs cross the neurovascular unit *in vitro* and *in vivo*

To validate whether PEI-MSNs could in principle be used in future for targeting GSCs *in vivo*, we screened their permeability through an *in vitro* model of BBTB[37]. Briefly, this model establishes a mimic of BBB by co-culturing mouse brain microvascular endothelial cells and astrocytes in Transwell inserts. Once the endothelial cells formed a tight monolayer, inserts were placed on BT-12 gliospheres and 100 ng of PEI-MSNs were added on the endothelial side. Passage of the PEI-MSNs through the inserts was followed by confocal microscopy **(Fig. S9)**. Lysotracker fluorescent dye was used to label the lysosomes. After 24h, PEI-MSNs were still detected in the endothelial cells and astrocytes and co-localized with lysosomes **(Fig. 6A-B)**. Interestingly, PEI-MSNs were also abundantly detected on the other side of the BBTB, in the lysosomes of the BT-12 gliospheres **(Fig. 6C)**. After 3 days, endothelial cells, astrocytes, and BT-12 cells were removed from the Transwells and their viability was measured by MTT. BT-12 gliosphere viability was 31% lower compared to untreated BT-12 cells isolated from the control BBTB **(Fig. 6D)**. The viability of endothelial cells and astrocytes was not significantly affected compared to the untreated cells (−3% and −6%, respectively) suggesting specific toxicity towards the BT-12 GSCs.

**Figure 6.**
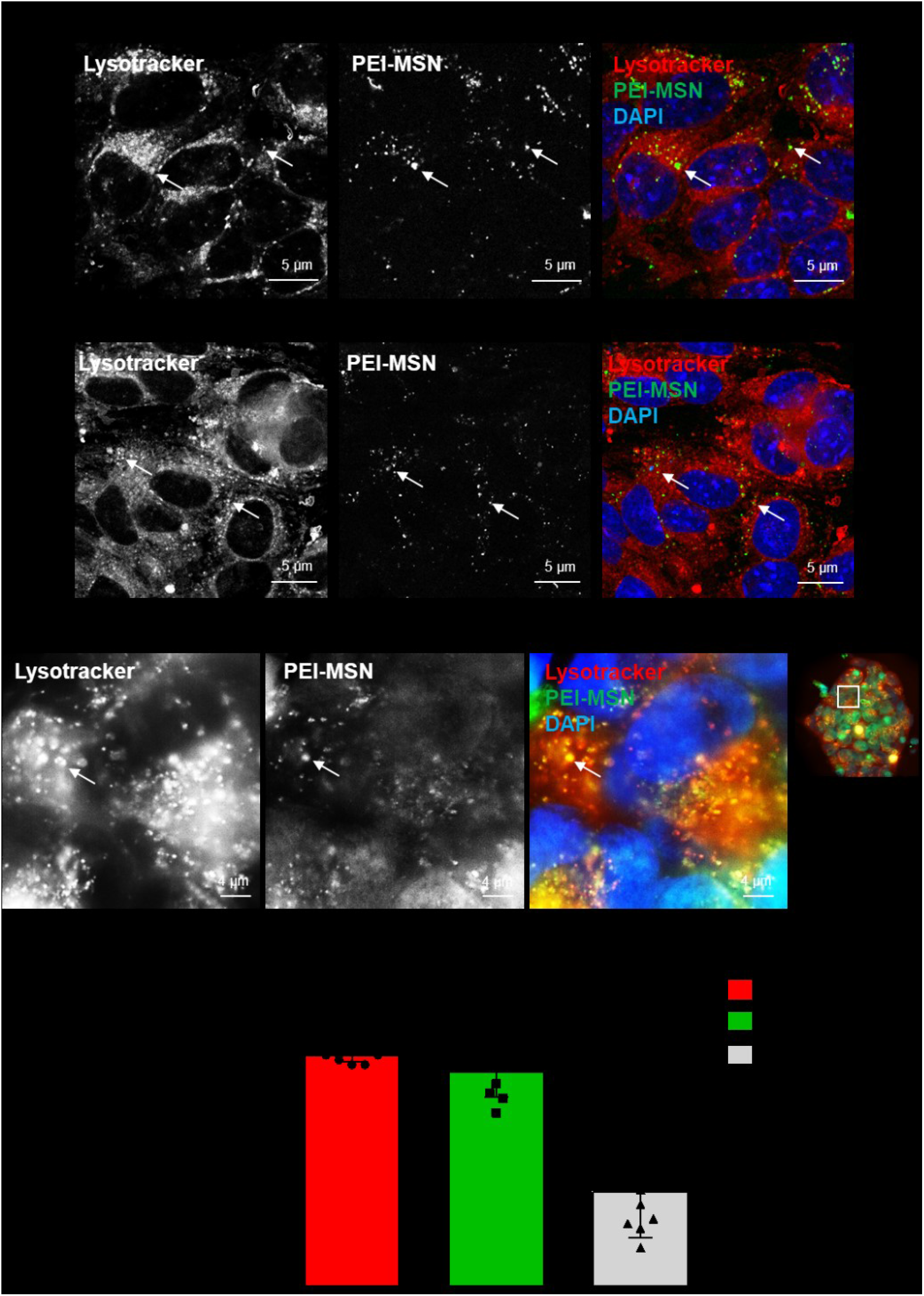
The PEI-MSNs cross the BBTB *in vitro*. A-C) Confocal images of the colocalization of PEI-MSN (green) with the Lysotracker dye (red) in the endothelial (A), astrocytic (B) and glioblastoma GSC (C) compartments 24h after addition of the PEI-MSN. Images in C are magnified from a BT-12 gliospheres (right panel). D) Cell viability measured by MTT after 72h. Values are normalized to control conditions (cells isolated from untreated BBTB). n = 3, three pooled experiments. P-value calculated with one-way ANOVA with Sidak’s multiple comparison test.

We then proceeded with the *in vivo* evaluation of the PEI-MSN passage through the neurovascular unit and brain distribution in immunocompromised mice implanted with BT-12 or U87MG-GFP cells. Due to the relatively small diameter (50 nm) of the PEI-MSNs, we evaluated two different administration routes, *i.e.* the classical caudal vein route or intranasal dosage [63]. Intranasal delivery of PEI-MSNs (35 μg in 5 μL of PBS) was distributed drop by drop, alternating between the nostrils. The procedure was repeated 3 times every two hours. Animals were euthanized 2 hours after completing the intranasal dosage and eight hours after IV injections of the PEI-MSNs, and brains were collected. We then verified the distribution of the particles in different areas of the brain parenchyma and the glioblastoma xenografts. We observed PEI-MSNs associated with or outside brain blood capillaries in different regions of the cerebral cortex following IV injections **(Fig. 7A & C)**. For the intranasally administered mice, no preferential distribution of the PEI-MSNs was observed around blood vessels, but nanoparticles could still be detected within various regions of the brain parenchyma, including posterior parts of the encephalon such as the hippocampus **(Fig. 7B & C)**. The intranasal administration rationale is based on the fenestrated BBB before the cribriform plate, allowing intracranial accessibility through the olfactory neuron endings. Compounds are endocytosed by the cilia, can travel inside the axons, and reach the central nervous system from the olfactory bulb. We then verified the distribution of the PEI-MSNs in the olfactory bulb of both IV and intranasally treated animals. We could observe a very high density of nanoparticles in the olfactory bulb of the intranasally administered mice **(Fig. 7D)**, supporting the brain accessibility of the PEI-MSNs through the vomeronasal nerves. As PEI-MSNs can also cross the BBB when delivered intravenously, we could detect PEI-MSNs in the olfactory bulb tissue of IV injected mice in similar densities than observed in the rest of the brain **(Fig. 7D)**. We eventually verified the presence of nanoparticles in the BT-12 and U87MG-GFP intracranial tumors **(Fig. 7E-F)**. Interestingly, PEI-MSNs could be observed in tumors after both IV and intranasal administration, although the IV injected animals seemed to exhibit a better intratumoral distribution with more NPs observed within the tumor tissue compared to the intranasal delivery **(Fig 7E-F)**.

**Figure 7.**
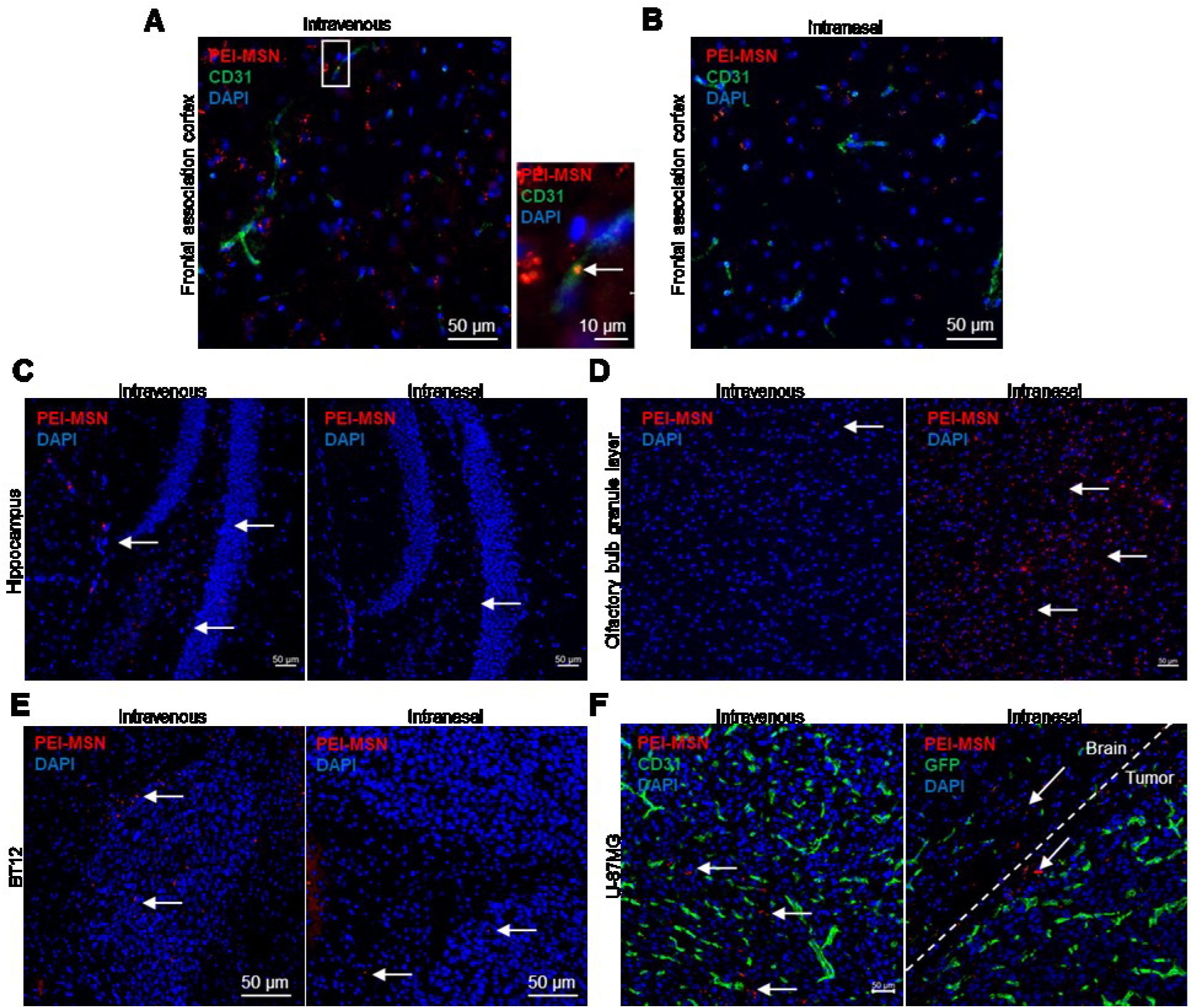
The PEI-MSNs can be delivered to the brain *in vivo*. A-B) Confocal images of the colocalization of PEI-MSN (red) with the endothelial cell marker CD31 (green) in the frontal association cortex of mice injected with 100 μg of PEI-MSNs in the caudal vein (A), or with 3 dosages of 35 μg of PEI-MSNs given intranasally (B). After 8h, PEI-MSNs were found associated with the brain endothelium (A, arrow right panel) and in the brain parenchyma. C) Confocal images of the PEI-MSNs (arrows, red) in the hippocampus area of IV-injected (left) or intranasally administered mice (right). D) Confocal images of the PEI-MSNs (arrows, red) in the olfactory bulb granular layer of IV-injected (left) or intranasally administered mice (right). E) Confocal images of the PEI-MSNs (arrows, red) IV-injected (left) or intranasally administered (right) mice xenografted with BT-12 cells. Cell nuclei are counterstained with DAPI. F) Confocal images of the PEI-MSNs (arrows, red) IV-injected (left) or intranasally administered (right) mice xenografted with U-87MG-GFP cells. Cell nuclei are counterstained with DAPI and blood vessels labeled with an anti-CD31 (green).

Intranasal delivery showed heterogenous distribution of the PEI-MSNs in the brain, *i.e.* very high concentration at the entrance point in the olfactory bulb and only a few PEI-MSNs in distant structures such as the hippocampus and the tumor. However, this provides a proof-of-principle that the central nervous system can be reached through this non-invasive method. Also, as the travel of the nanoparticles endocytosed at the nerve endings is certainly slower than the blood flow, longer timepoints could show an enhanced PEI-MSN distribution in the brain tissue.

## Conclusions

Our results indicate that a PEI surface coating on MSNs can specifically and effectively induce death of GSCs, whereas other cancer cell lines are not affected. This finding contradicts the common consensus that GSCs are considered to be the most resistant cell population in the tumor, supporting the existence of undiscovered vulnerabilities in GSC biology. From a mechanistic point of view, the results show that PEI-MSNs accumulate in the lysosomes of BT-12 cells after cellular uptake. This, in turn, leads to the “proton sponge effect”, causing widespread localization of PEI-MSNs and lysosomal enzymes such as cathepsins and other hydrolases, into the cytoplasmic space via lysosomal membrane rupture. Release of the lysosomal enzymes, in turn, leads to cell death. Similar vulnerability in the GSCs was recently found to be induced by antihistaminergic drugs[15]. Therefore, our data confirm this finding and strengthens the theory of the lysosomal vulnerability of GSCs. Our results further suggest that PEI modification imparts a new, inherent property to nanoparticles in selective killing of GSCs without any additional anti-cancer drug treatment, which is contrary to recently reported drug based nanomedicinal approaches to eradicate GB [64–67]. We also demonstrated the potential of the PEI-MSNs to be directly delivered to GSCs via intranasal or intravenous administration for the successful eradication of GSCs *in vivo*. Taken together, these results support the importance of discovering novel vulnerabilities in lysosome-associated pathways, GSCs, and in this specific case; the hidden potential in the inherent activity of potential drug delivery systems. In addition, this study implies compelling evidence for the therapeutic application of the proton-sponge effect by cationically surface modified nanoparticles to target cells with vulnerable lysosomes.

## Author contribution

All authors have approved the final version of the manuscript. N.P. designed the study, performed live-cell microscopy, and wrote the manuscript. J.M. performed the colony growth assay and western blotting and wrote the manuscript. V.L.J. performed the *in vivo* experiments, analyses using the *in vitro* BBB model, and wrote the manuscript. M.P. performed the TEM experiments. D.S.K. and E.C. performed particle synthesis and characterization. P.L., J.W., and J.R. supervised the study, provided guidelines, and edited the manuscript.

## Acknowledgments

The authors are very grateful to Jenni Laine at the Electron Microscopy unit, the University of Turku for sample processing. Jane and Aatos Erkko Foundation, Academy of Finland (project #309374) and Sigrid Jusélius Foundation are acknowledged for funding.

## Supplementary information

**Figure S1.**
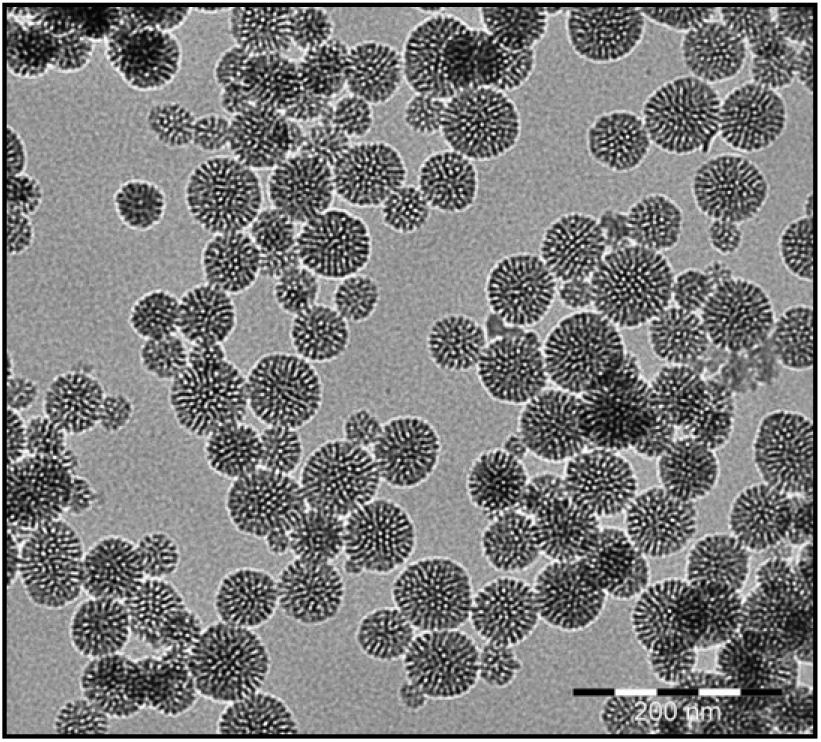
TEM micrographs of the spherical PEI-MSNs particles of an approximate size of 50 nm with porous structure. Scale bar = 200 nm.

**Figure S2.**
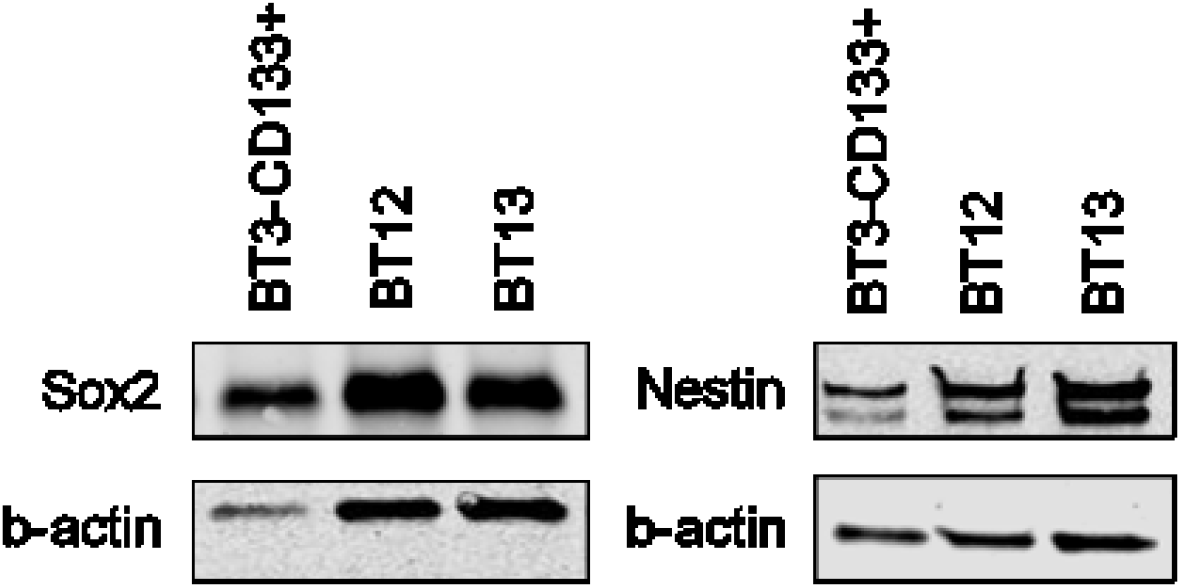
Validation of GSCs (BT-3-CD133^+^, BT-12, and BT-13) using Sox2 and nestin (stem cell markers).

**Figure S3.**
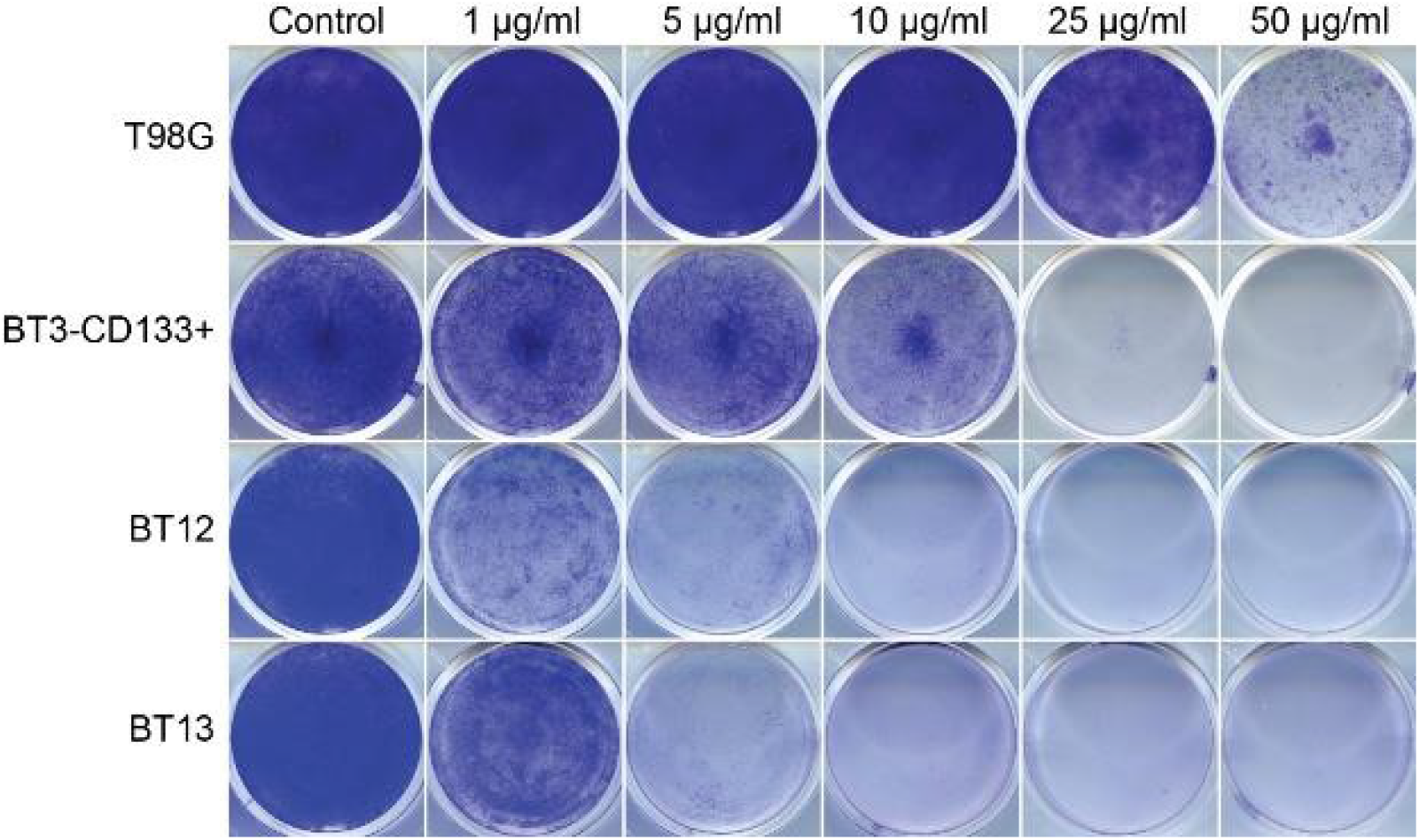
Selective cell death of patient-derived GSCs by PEI-MSNs. A) Colony growth assay analysis by using crystal violet staining of T98G (GB cell line), BT-3-CD133^+^, BT-12 and BT-13 GSCs treated with 1-50 μg/mL of **PEI-MSNs**.

**Figure S4.**
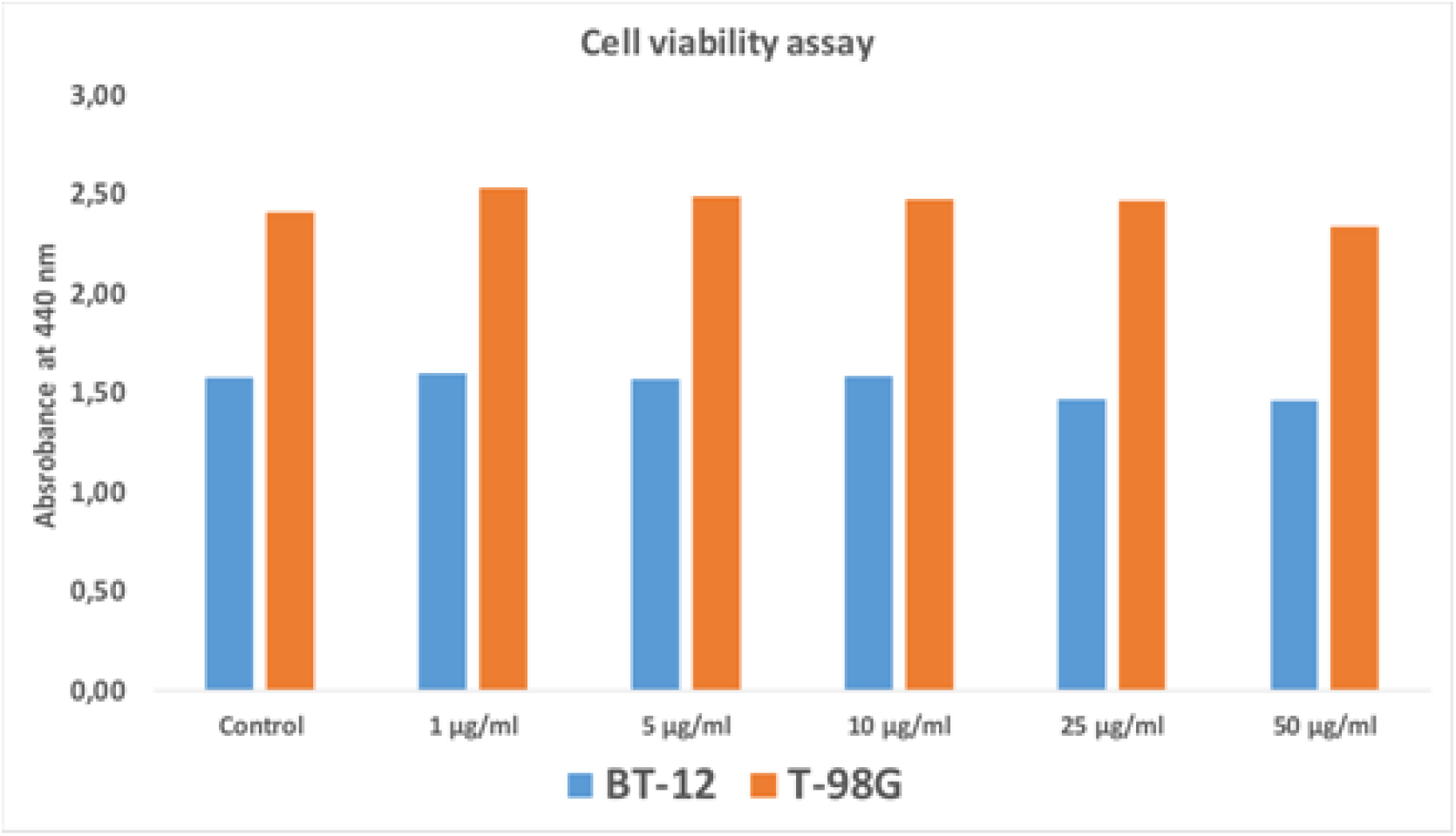
Viability of BT-12 GSCs and T98G (GB) cells treated with 1-50 μg/mL of **plain MSNs (without PEI)**. The cell viability was assessed by WST-1 cell proliferation assay.

**Figure S5.**
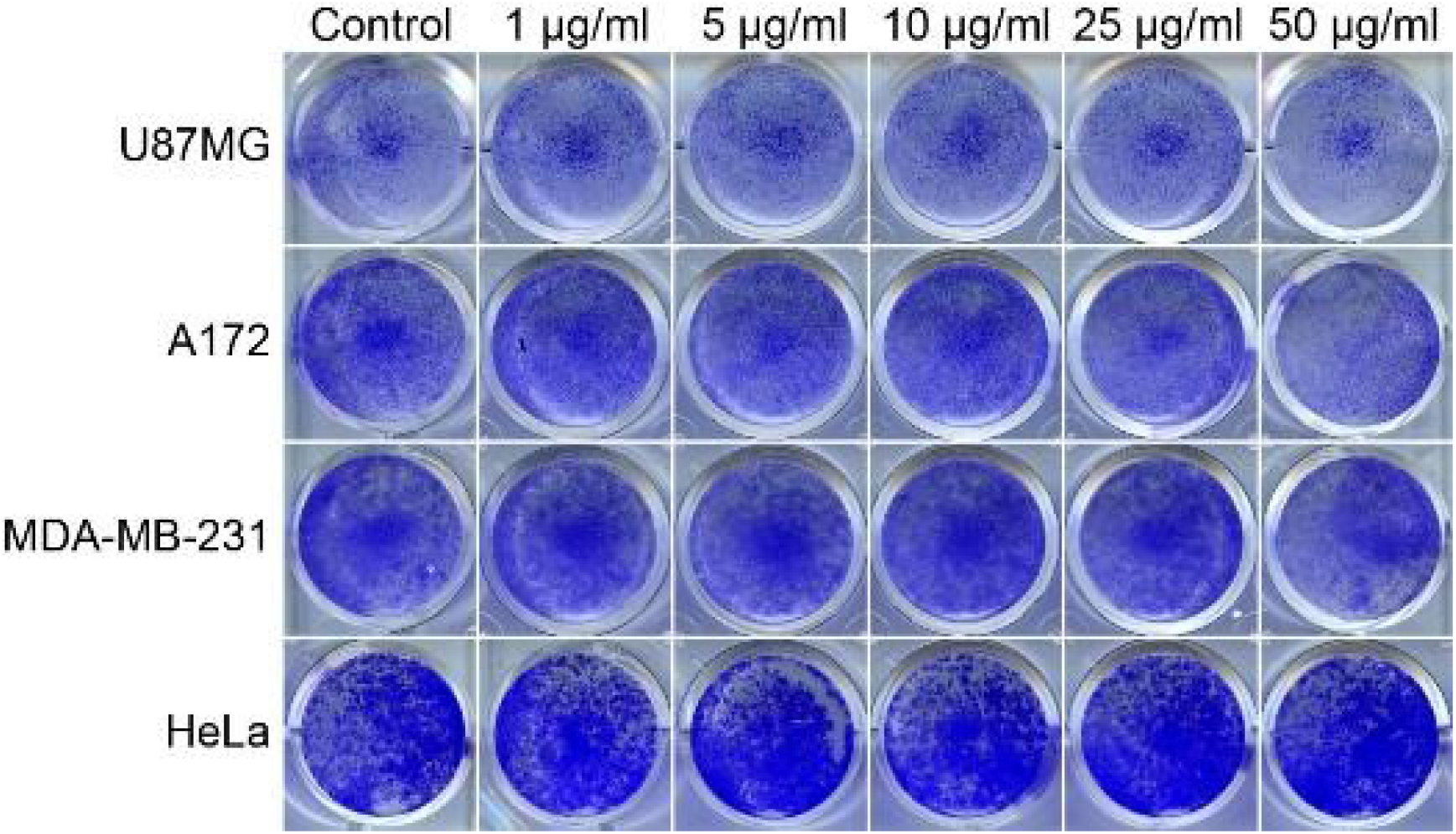
Colony growth of U87MG (human GB cells), A172 (human GB cells), MDA-MB-231(human breast cancer cells) and HeLa cells (human cervical cancer cells) treated with 1-50 μg/mL of **PEI-MSNs**.

**Figure S6.**
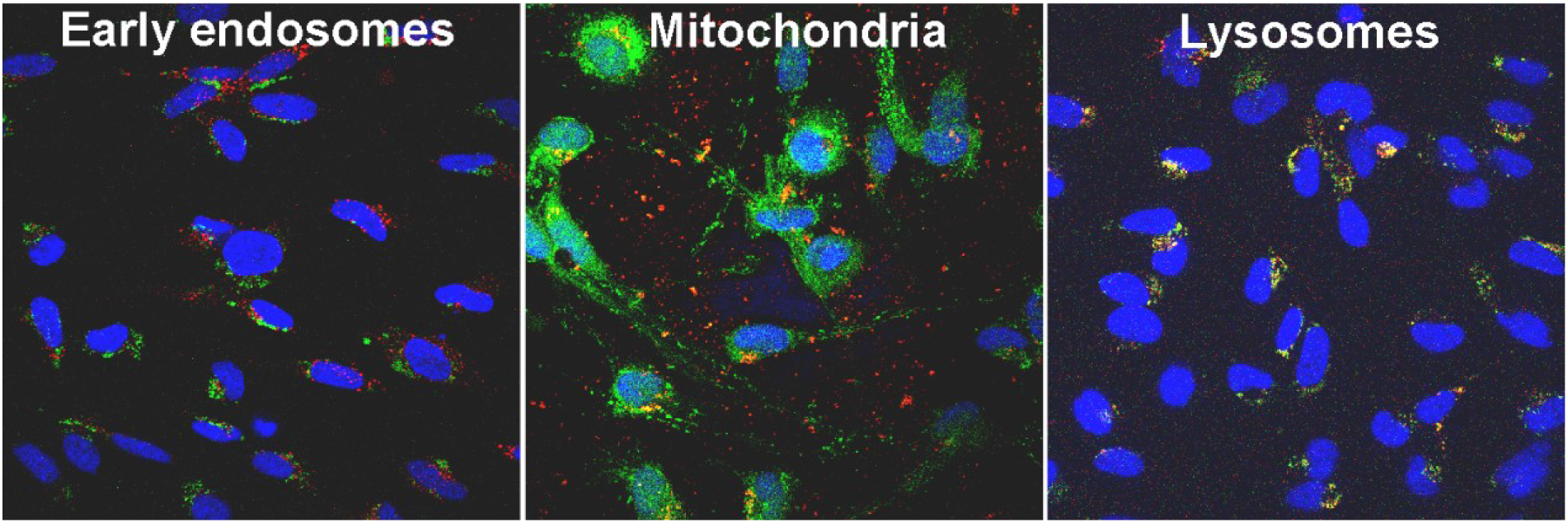
Localization of the PEI-MNSs in the treated BT-12 GSCs by confocal microscopy. Intracellular localization of PEI-MSNs (red) was studied using markers of early endosomes (EEA-1), mitochondria (Mitotracker), and lysosomes (LAMP-1) (green color). The nuclei were visualized by using DAPI (blue). Co-localization is seen in yellow color.

**Figure S7.**
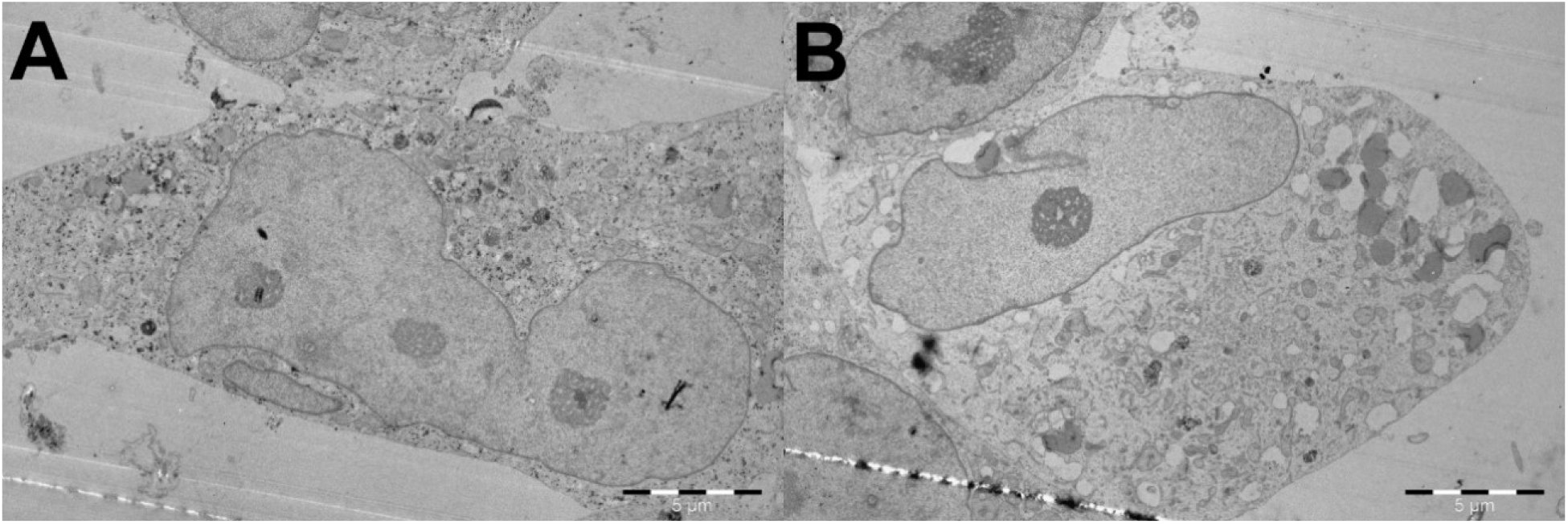
TEM images of PEI-MSNs treated BT-12 GSCs. No PEI-MSNs were detected within the nuclear space.

**Figure S8.**
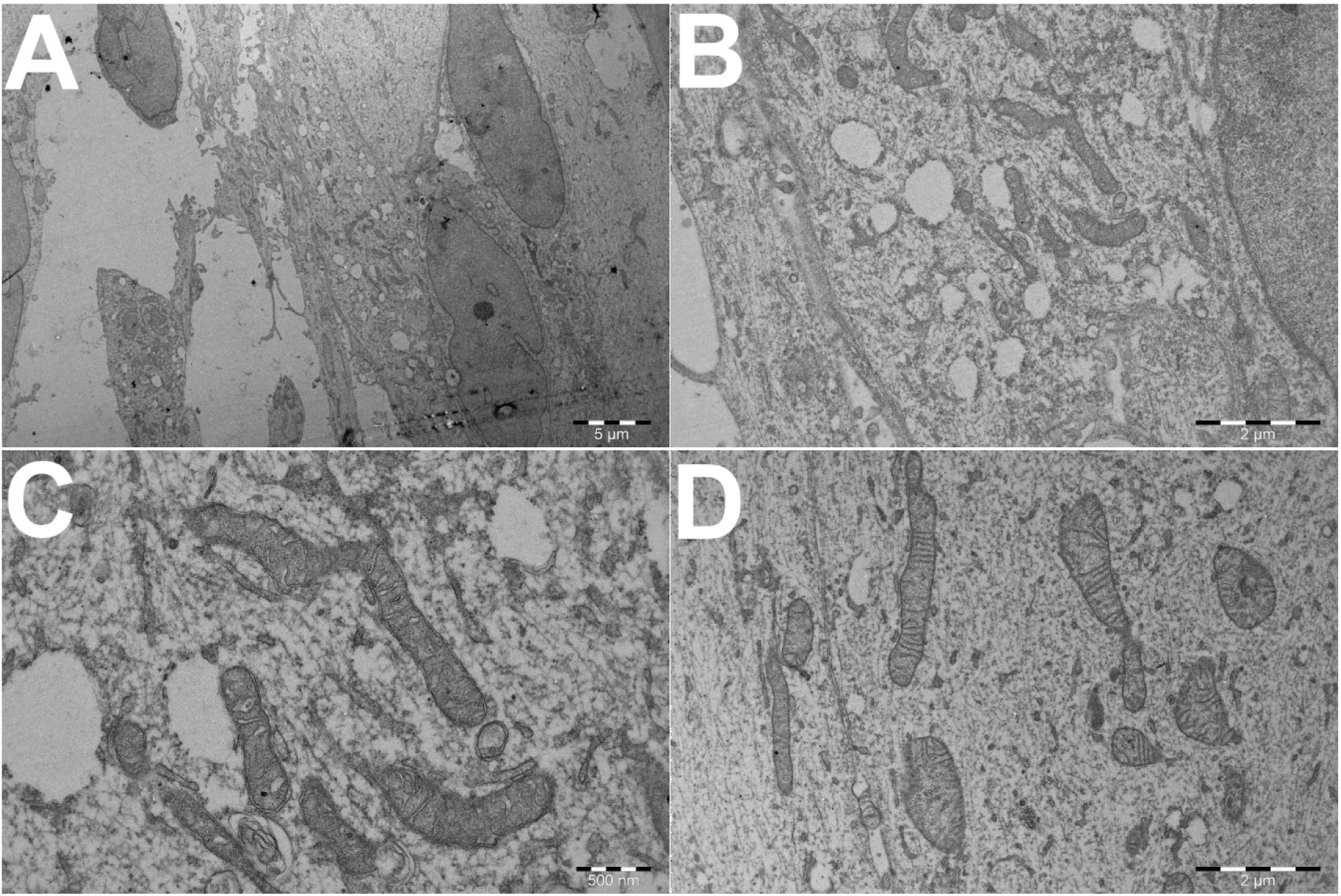
TEM images of control BT-12 GSCs without PEI-MSNs treatment. A) Overview of a cell. B) Empty vesicles and normal ultrastructure of BT-12 GSCs. C-D). A close overview of the intact mitochondrial structure.

**Figure S9.**
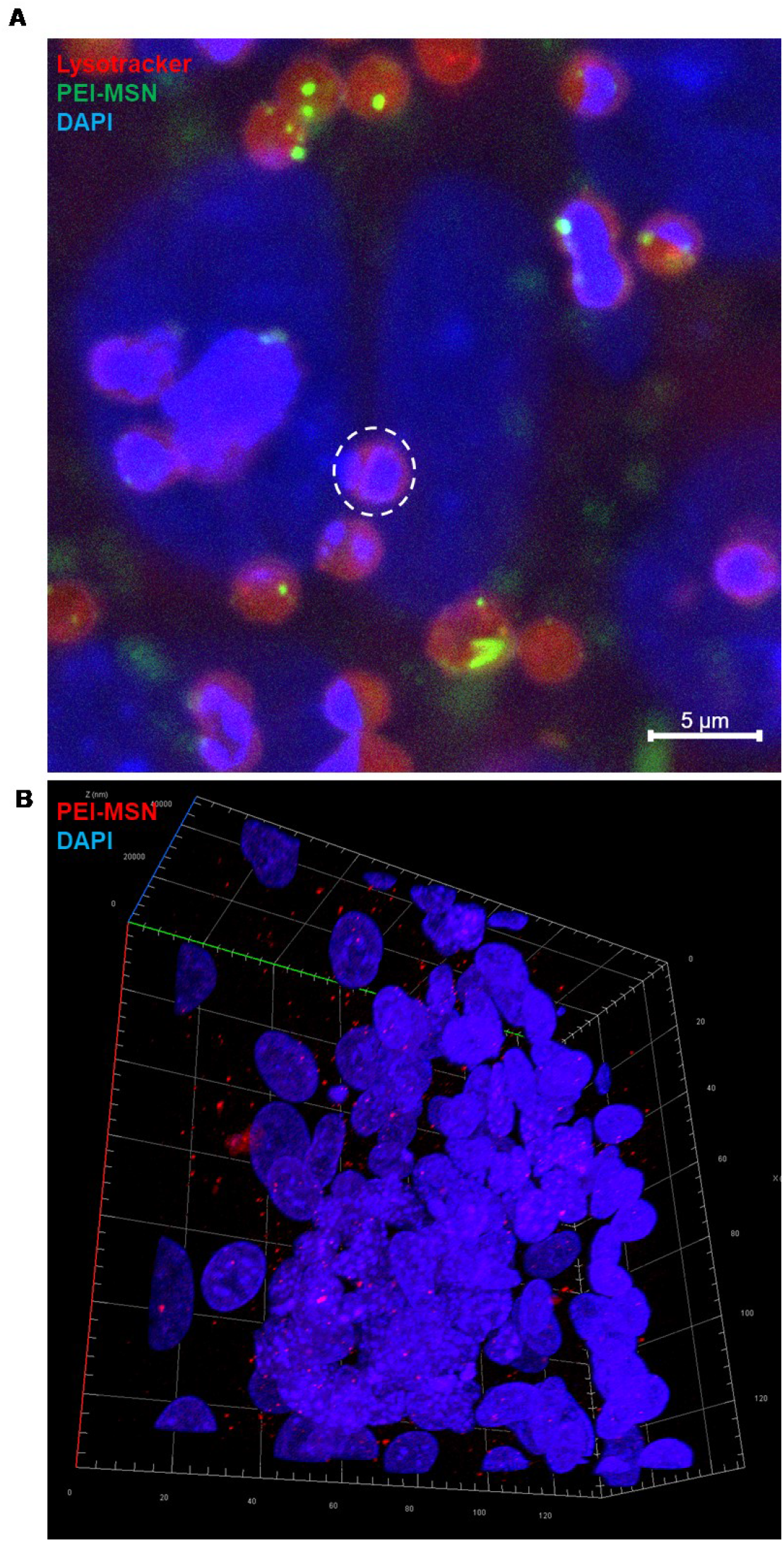
A) Sagittal confocal view of the Transwell membrane (with pores highlighted by a dotted circle) showing PEI-MSN (green) transcytosis through the endothelial cells (Lysotracker red and nuclei blue). B) 3D reconstruction from confocal Z-stacks of a BT-12 gliosphere isolated from the BBTB. PEI-MSNs (red) are distributed all around and penetrated inside the sphere.

